# Zebrafish as a model to investigate the effects of exercise in cancer

**DOI:** 10.1101/279232

**Authors:** Alexandra Yin, Nathaniel R. Campbell, Lee W. Jones, Richard M. White

## Abstract

Emerging data indicates that exercise may regulate cancer pathogenesis, but the mechanisms underpinning how it regulates the tumor as well as surrounding microenvironment are poorly understood. Dissecting this complex, highly integrated physiology requires model systems which accurately recapitulate key aspects of human response to exercise, yet permit rapid and unbiased genetic interrogation of relevant pathways. The zebrafish has emerged as a new model for cancer due to its high resolution in vivo imaging and capacity for large-scale, unbiased screening approaches. Here, we have developed a set of tools to study the effects of exercise in a zebrafish model of melanoma. Using a flow chamber, we studied the effects of endurance exercise bouts (3-6 hour/d, 5d/wk for 1 to 3 wks) in both larval and adult zebrafish. The regimen was well tolerated, with no unexpected toxicities or changes in survival. When the zebrafish were transplanted with ZMEL1-GFP melanoma cells, we found that endurance exercise over a 2-week period led to a significant decrease in cancer growth in the larval zebrafish. As zebrafish cancer models show strong conservation in human disease, our findings have direct application to understanding the human exercise/cancer relationship.

## Introduction

Observational evidence links regular exercise to a reduced primary incidence of several forms of cancer as well as a reduced risk of relapse and cancer mortality after diagnosis (Friedenreich et al., 2016). In support, emerging data demonstrates that exercise has antitumor activity in some, but not all, allograft or clinically derived xenograft rodent models of cancer (Ashcraft et al., 2016). The mechanisms underpinning how exercise regulates the tumor as well as surrounding microenvironment are poorly understood (Koelwyn et al., 2017). Dissecting this complex, highly integrated physiology requires model systems which accurately recapitulate key aspects of human response to exercise, yet permit rapid and unbiased genetic interrogation of relevant pathways. While the majority of preclinical investigations in exercise-oncology have been conducted in rodent models (Ashcraft et al., 2016), limited work to date has utilized “lower” metazoan species such as worms (Caenorhabditis elegans), flies (Drosophila melanogaster), and zebrafish (Danio rerio) (Hawley et al., 2014). These models offer several distinct features that may be particularly advantageous for investigation of exercise-disease mechanisms.

Zebrafish are a small vertebrate model organisms with several novel characteristics ideal for investigating exercise regulation of cancer. First, fish are highly amenable to cancer modeling of a wide variety of tumors, ranging from melanoma to rhabdomyosarcoma amongst others (White et al., 2013). Second, zebrafish are uniquely suited to high throughput interrogation and genetic perturbation of both tumor cells as well as those in the surrounding microenvironment. Third, these models are readily applicable to unbiased screens using small molecules or CRISPRs that allow for the discovery of unexpected pathways that might be relevant for exercise oncology. Fourth, the availability of optically transparent strains such as *casper* allow for unprecedented imaging of the tumor/microenvironment interface at single-cell resolution (White et al., 2008). Finally, the physiology of the zebrafish closely recapitulates many key vertebrate aspects of endurance exercise - fish habitually swim throughout the day at low speed, but when challenged with a high water flow environment markedly increase swimming speed through sustained skeletal muscle contraction. The fish have an advanced circulatory system permitting appropriate cardiovascular response to endurance exercise-induced increased metabolic demand.

Our laboratory utilizes a well-established transgenic zebrafish model of melanoma to examine questions with relevance to the human disease. In these models, human oncogenic BRAF^V600E^ is transgenically expressed under the melanocyte-specific mitfa promoter (Patton et al., 2005). When crossed with p53 mutant animals, the resultant animals develop a 100% penetrant melanoma that strongly resembles the human disease at histologic, gene expression and genomic levels (White et al., 2011; Kaufman et al., 2016; Ceol et al., 2011). More recently, this model has been extended to the study of metastatic disease. A zebrafish melanoma cell line called ZMEL1 was derived from the BRAF^V600E^ transgenic animals, and when transplanted into the *casper* recipient, metastatic disease can be imaged at very high resolution over time (Heilmann et al., 2015). The ZMEL1 model has been further utilized to examine the role of the microenvironment in metastasis (Kim et al., 2017), in which CRISPR/Cas9 approaches can be used to delete genes such as EDN3b or ECE2b from the stroma.

In this study, we leverage these zebrafish melanoma models to understand the effect of endurance exercise on melanoma growth. In this proof-of-principle methodology study, we have developed a set of tools to create and accurately quantify a system for endurance exercise in this model, and then leverage these tools to demonstrate, for the first time, that exercise strongly suppresses tumor growth. We contend that this system provides a new paradigm for a wide variety of future studies aimed at dissecting the mechanisms of exercise-oncology in an unbiased manner.

## Results

### Establishment of an endurance exercise system for zebrafish cancer models

The zebrafish has been widely adopted as a cancer model (White et al., 2013). This is most readily observed in melanoma, with transgenic (Patton et al., 2005; White et al., 2011; Ceol et al., 2011) and transplantation methods (Heilmann et al., 2015) established to interrogate various aspects of melanoma biology. Indeed, zebrafish models of melanoma are particularly advantageous since they permit elucidation of the tumor cell-intrinsic and microenvironmental features that mediate tumorigenesis and metastasis. Such aspects are of direct relevance in the present context since exercise-induced changes in whole-organismal physiology likely exert antitumor effects via modulation of the tissue microenvironment/landscape - cancer cell interaction. Our goal was to leverage these aspects of melanoma models to build a system for exercise in the zebrafish.

As such, the first objective was to establish a reproducible paradigm of endurance exercise in zebrafish. Previous work has established a flow chamber systems (Blazka-type or Loligo chamber) in which water rapidly flows past the zebrafish which elicits an innate upstream swim response (Gilbert et al., 2014; Palstra et al., 2010). An increase in flow rate beyond normal background water flow (in fish tanks) requires voluntary skeletal muscle activation to maintain the swim response, which is a paradigm of exercise (i.e., any bodily movement produced by skeletal muscles that requires energy expenditure). Because the flow rate can be controlled, this allows for precise implementation as well as quantification of the exercise dose / volume.

The swim chamber applied in this study is shown in Figure 1. In brief, the apparatus consists of a flow chamber with fixed water volume connected to a controllable pump that regulates water flow rate. This is critical since this permits precise quantification of flow rate (i.e., exercise intensity) as well as investigation of dose/exposure - response relationships. Adult fish are directly placed into the water chamber allowing visual observation during rest and exercise periods.

**Figure 1:**
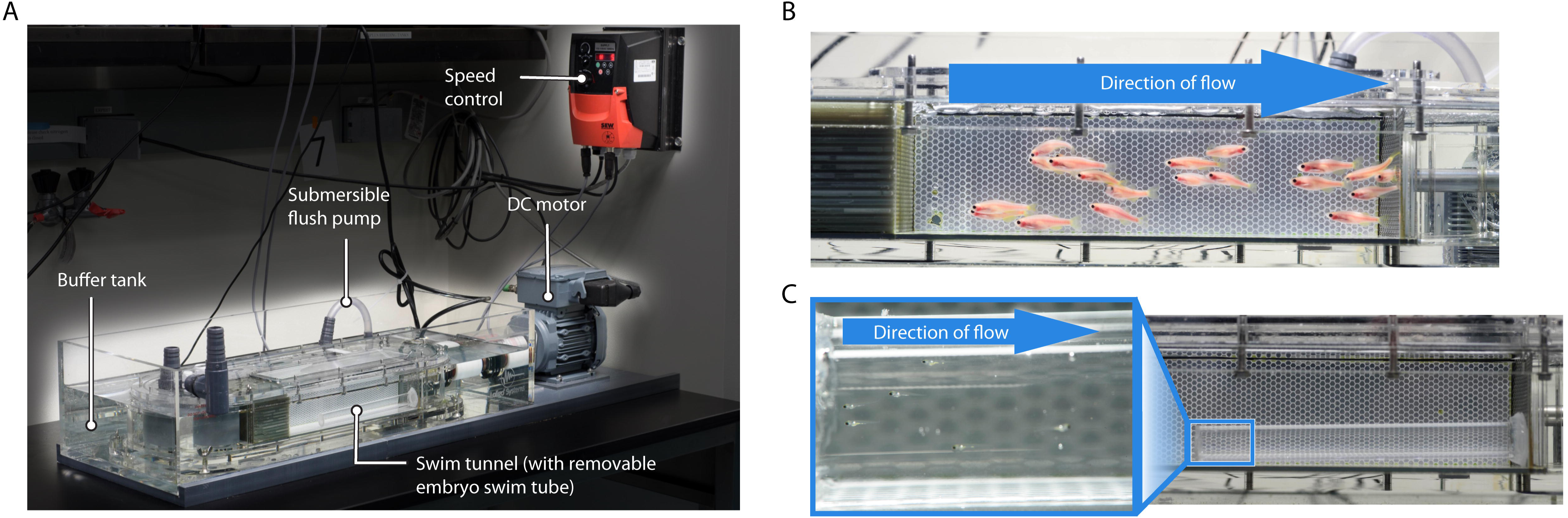
An exercise chamber for zebrafish. (A) The overall design of the Loligo swim tunnel system. (B) Appearance of the adult zebrafish within the main swim tunnel chamber. (C) Appearance of the larval zebrafish within the ZEST chamber, with higher magnification shown to the left.

We also adapted this system for younger fish, in the embryonic/larval period. This has advantages because many of the transplantable cancer assays have been optimized for this larval period (Fior et al., 2017). In addition, because fish are smaller in the embryonic/larval stage, it allows for the investigation of a much larger number of fish. However, due to their small size, the existing Loligo chamber was not appropriate, as the larvae would be pulled through the downstream grate, and therefore not confined to the swim tunnel. An exercise tunnel was built to adapt the 5L Loligo swim tunnel to exercise the embryonic/larval zebrafish, which are around 4-5mm in size when they start exercise 12 days post fertilization. We termed this embryonic adaptation the ZEST, for “Zebrafish Embryo Swim Tunnel”. One end of an acrylic tube was covered in fine mesh, which was adhered to the acrylic tube using silicon glue to ensure that the larval zebrafish were confined to the swim tunnel. A cap was applied to the opposite end of the ZEST using a rubber tube with a diameter slightly larger than the outside of the acrylic tube. The end of the cap was covered in the same fine mesh, which was also adhered to the cap using silicon glue. Before putting the larvae into the ZEST, the unit was submerged in system water within the swim tunnel. Larvae were placed into the ZEST using a transfer pipette, and the cap was added to ensure that the larvae remained within the ZEST. The ZEST was then located at the bottom of the existing Loligo swim tunnel. These systems provide the capacity to conduct exercise experiments in virtually any age fish, from 2 days post fertilization to well over 1 year post fertilization.

### Exercising zebrafish exhibit characteristic “drafting” behaviors

To understand how the zebrafish adapted to the exercise system, we used time-lapse imaging of an acute exercise bout in adult zebrafish. As shown in Figure 2 and Supplemental Movie 1, one of the characteristic behaviors we noted is best described as “drafting” in which one fish acts as the leader of the pack, closely followed by a group of fish. After several seconds, we observed the leader fish falling to the rear of the pack, replaced by a new fish at the group front. All fish then resume upstream swimming behavior, and the cycle is repeated continuously for the duration of the exercise bout. This data suggests that zebrafish, not unlike rapidly running mammals, take advantage of nearby fish to draft, therefore enhancing exercise efficiency. It also suggested that over time, individual fish are likely working at submaximal levels, allowing new fish to lead. This indicated the need to perform formal “exercise tolerance” testing to determine endurance exercise capacity thereby permitting appropriate exercise regimen dosing in experiments evaluating the effects of chronic endurance exercise.

**Figure 2:**
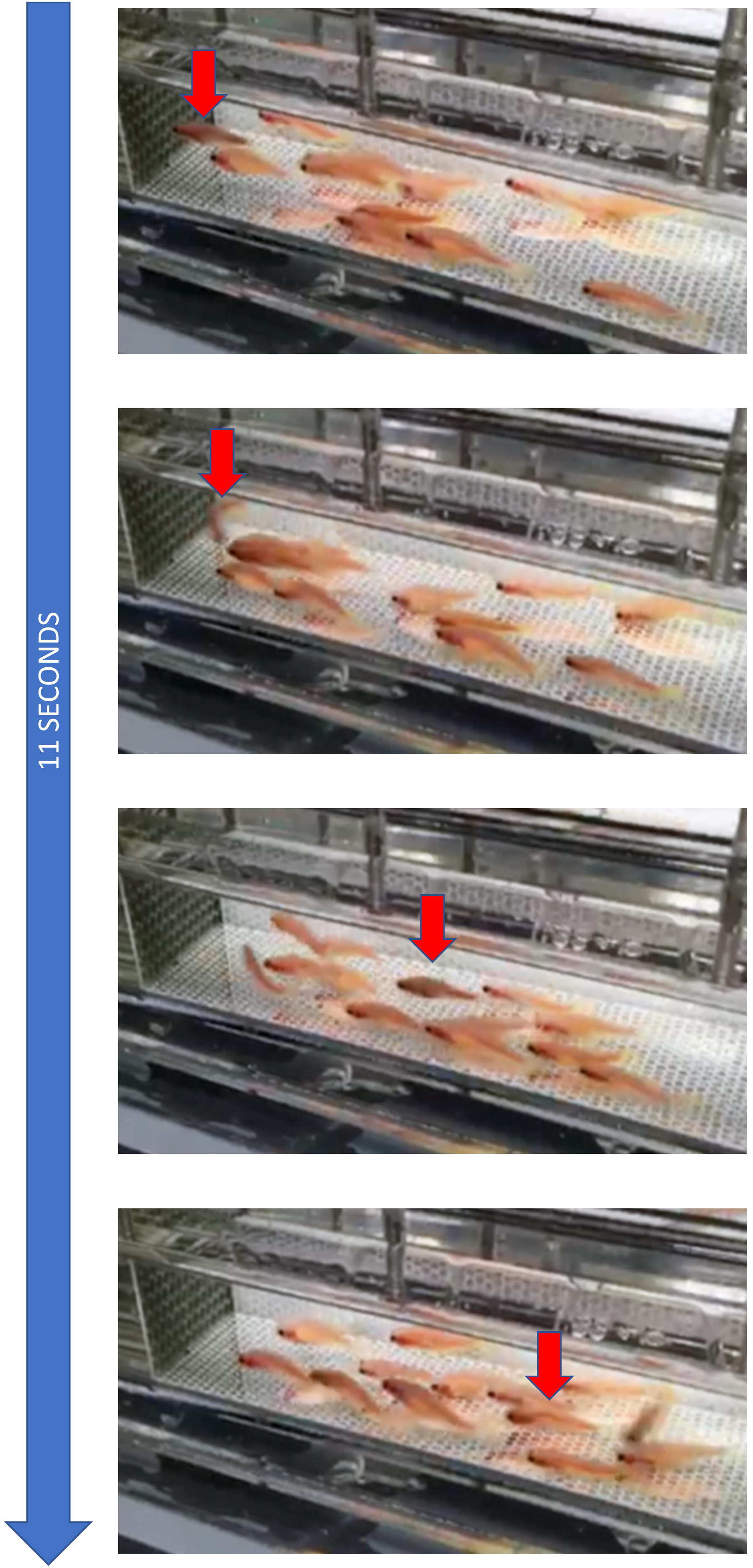
A time lapse movie of adult zebrafish swimming at maximal capacity. (A) The leader fish is marked with a red arrow, with a fish just behind it “drafting” off of it. (B) Several seconds later, this leader fish flips around 180 degrees and begins to fall backwards. (C, D) The position of the former leader fish is now seen to continue to fall away towards the back of the pack.

### Larval Zebrafish Exercise Regimen

Dosing of endurance exercise for the larval zebrafish was determined using a speed test. 12 days post fertilization (10 days post transplant), groups of ∼50 larval fish were allowed to acclimate to the ZEST for 5 minutes. The water flow speed was initially set at 5 cm/s then increased 1 cm/s every 2 minutes. In the event that a fish remained at the back grate before the speed of 10 cm/s for greater than 5 seconds, the fish was removed from the group and considered a “non-swimmer”. As the speed increased, we observed that many fish intermittently clung to the side of the tube in order to maintain their position within the ZEST. Speed was increased to a maximum of 10 cm/sec, at which point >50% of the larvae remained at the back grate, which we considered the maximal dose. Throughout the two-week experimental period, the frequency at which the larvae clung to the side of the tube decreased, suggesting a training adaptation. At a speed of 8 cm/sec, the majority of the larvae maintained position in the ZEST, not requiring clinging behavior. In subsequent experiments, a speed of 8 cm/sec (i.e., ∼75% of maximal intensity) was tested as the intensity of endurance exercise.

### Adult Zebrafish Exercise Regimen

Groups of ∼30 adult fish were moved to the swim chamber of the 5L Loligo device and allowed to acclimate for 5 minutes. The speed of the water flow was set to 5 cm/s and the velocity was increased at increments of 2.5 cm/s at 2 minute intervals. Fish were observed throughout the test to ensure that no erratic swimming behavior developed. Any fish that could not maintain its position within the swim tunnel was caught by a mesh grate at the back of the tunnel. If a fish was pushed to the back grate before the speed of 10 cm/s was reached, and could not extract itself after 5 seconds, that fish was removed from the group and considered to be a “non-swimmer”. When the velocity of flow reached 30-35 cm/s, the fish swimming behavior started to become erratic and many of the fish were unable to maintain their position within the tunnel and were subsequently pressed against the downstream grate. As a result, a speed of 25 cm/s was chosen, which was about 75% of the determined maximal speed.

### Effects of Exercise on Tumor Progression

To test the effect of endurance exercise on melanoma growth, we used our previously developed zebrafish melanoma transplantation assay using the ZMEL1-GFP line (Heilmann et al., 2015; Kim et al., 2017). The overall experimental scheme is shown in Figure 3. The transgenic MiniCoopR system (Iyengar et al., 2012) was used to create a zebrafish melanoma, in which the mitfa promoter drives human BRAF^V600E^ in concert with a p53 loss of function allele, along with a GFP marker gene. From that original tumor, we created the ZMEL1-GFP cell line, which can be transplanted into transparent *casper* fish (White et al., 2008) to image tumor growth and metastasis (Heilmann et al., 2015). The cell line can be transplanted into either embryonic/larval *casper* recipients or adult *casper* recipients. There are advantages to either approach. When the *casper* recipient is a larvae, the ZMEL1-GFP cells grow and disseminate very rapidly. This is because these young fish have an innate, but not adaptive, immune system, and the larval environment provides a plethora of growth factors which accelerate tumor growth. However, these larval recipients may not recapitulate all aspects of the physiology seen in adults. In contrast, transplantation into the adult recipients has a fully intact physiology, but the downside is that the recipients require irradiation to prevent MHC mismatch between the ZMEL1-GFP cells and the individual casper recipient. Given the complexities of both models, both assays are utilized in investigating candidate therapeutic strategies.

**Figure 3:**
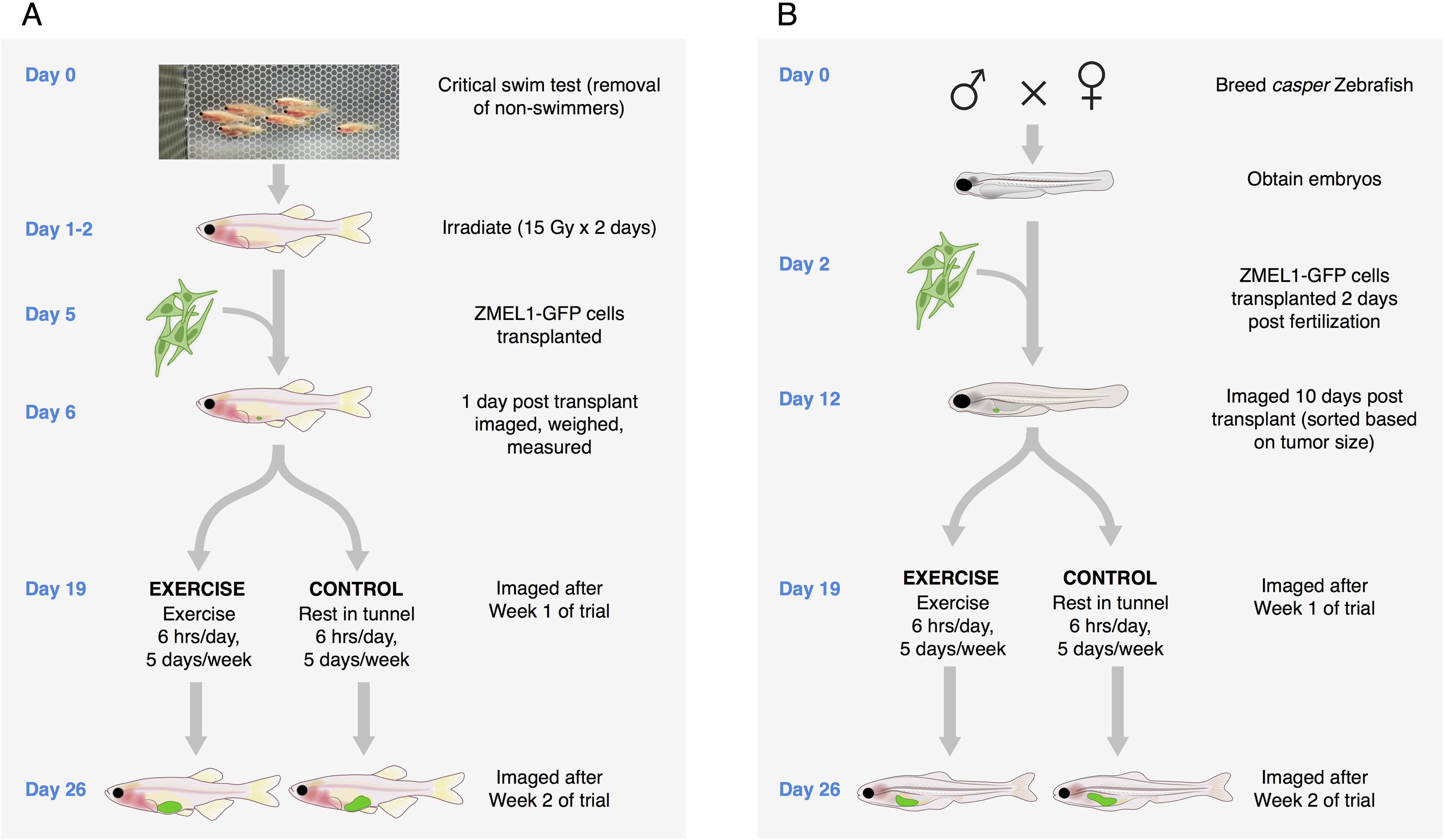
Experimental setup for testing the effects of exercise on melanoma. (A) Schema for the adult melanoma transplant assay. (B) Schema for the embryonic/larval melanoma transplant assay.

### The effect of exercise on melanoma using the embryonic/larval assay

We first examined the effect of exercise using the embryonic/larval ZMEL1-GFP melanoma assay (Figure 4). We transplanted ∼25 ZMEL1-GFP melanoma cells into the yolk sac of 2 day post fertilization zebrafish. The fish were imaged immediately after transplantation to ensure even distribution amongst the fish, and randomly allocated to endurance exercise or sham control. Endurance exercise consisted of swim training 6 hours/day for 5 consecutive days (Monday to Friday) with rest on weekends. Control groups were placed in the swim tunnel with no water flow for the same duration of time, but still otherwise transplanted in the same way. Each fish was individually imaged using brightfield, RFP and GFP fluorescence at 1 and 2 weeks post transplant. We also measured fish length as a proxy for overall growth, along with overall survival.

**Figure 4:**
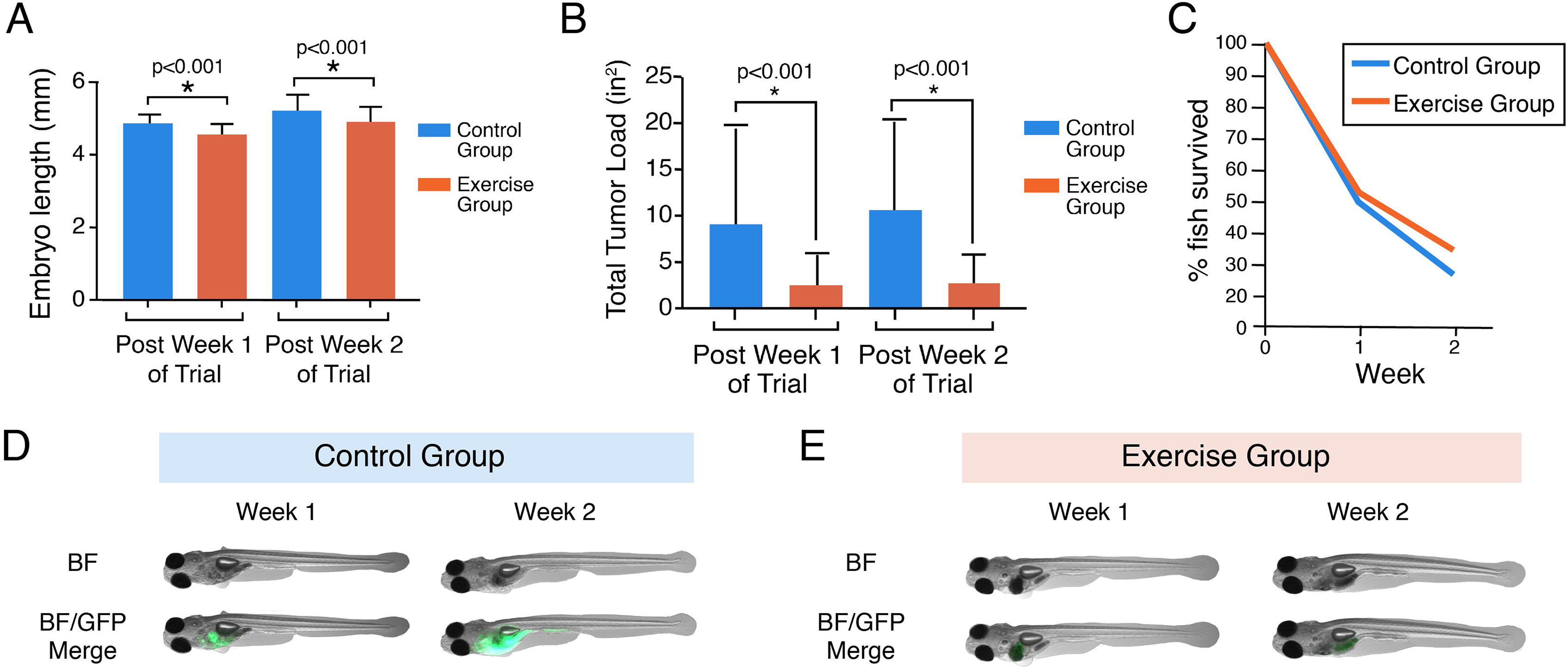
Larval exercise trial in melanoma. (A) Fish length over the period of the experiment showed no significant difference in control vs. exercise group. (B) Overall melanoma tumor size was assessed by measuring GFP+ area of the ZMEL1-GFP cells, after correcting for background autofluorescence. The exercised group showed a significant decrease in overall tumor size at both 1 and 2 weeks (*=p<0.05, Mann Whitney test). (C) Overall survival was similar between control and exercised fish. (D) Representative fish are shown at week 1 and 2 of the trial.

Images of the larvae were obtained at baseline and two weeks following endurance exercise. Tumor size was quantified using ImageJ. The background was subtracted using the Rhodamine channel to determine autofluorescence, and total GFP was quantified. Outliers were removed using the ROUT method of outlier identification. After both Week 1 and Week 2 of exercise, we found a significant 34% reduction in tumor size in the exercise group (n=28) compared to the control group (n=34) (p<0.001, Mann-Whitney test). This was accompanied by a small but not significant increase in survival in exercise versus control fish (35% versus 27%, respectively, p=0.28, Fischer’s Exact test). Fish length (extrapolated from images taken pre-trial and post-trial) was smaller in exercised versus control fish (4.57mm versus 4.90mm, respectively, p<0.001). Because the fish length is a relative indicator of growth during the larval period, this indicates that the intensity of exercise was sufficient to divert resources away from normal growth.

### The effect of exercise on melanoma using the adult assay

We next assessed the effect on melanoma growth using the adult transplant assay (Figure 5). Adult fish were anesthetized, weighed, and placed into one of four size categories (ie. small, medium, large, extra-large) based on their relative weight. Using weight as a stratification factor, animals were randomly allocated to exercise or sham control. All fish were imaged at baseline before randomization and at follow up two weeks later. Each fish was euthanized and imaged using brightfield, GFP, and Rhodamine filter sets and analyzed similarly to the larval assay. In contrast to the findings in the larval assay, no significant differences in the adult melanoma growth in control (n=122) versus exercise groups (n=118) were observed. There was also a small trend towards worse survival in the exercise group (60% in control versus 53% in exercise group). Consistent with this, we saw no differences in overall fish weight, suggesting that this dose of exercise may not have been sufficient to induce physiologic changes suggestive of weight loss, or that other factors precluded seeing an effect of exercise in this particular assay.

**Figure 5:**
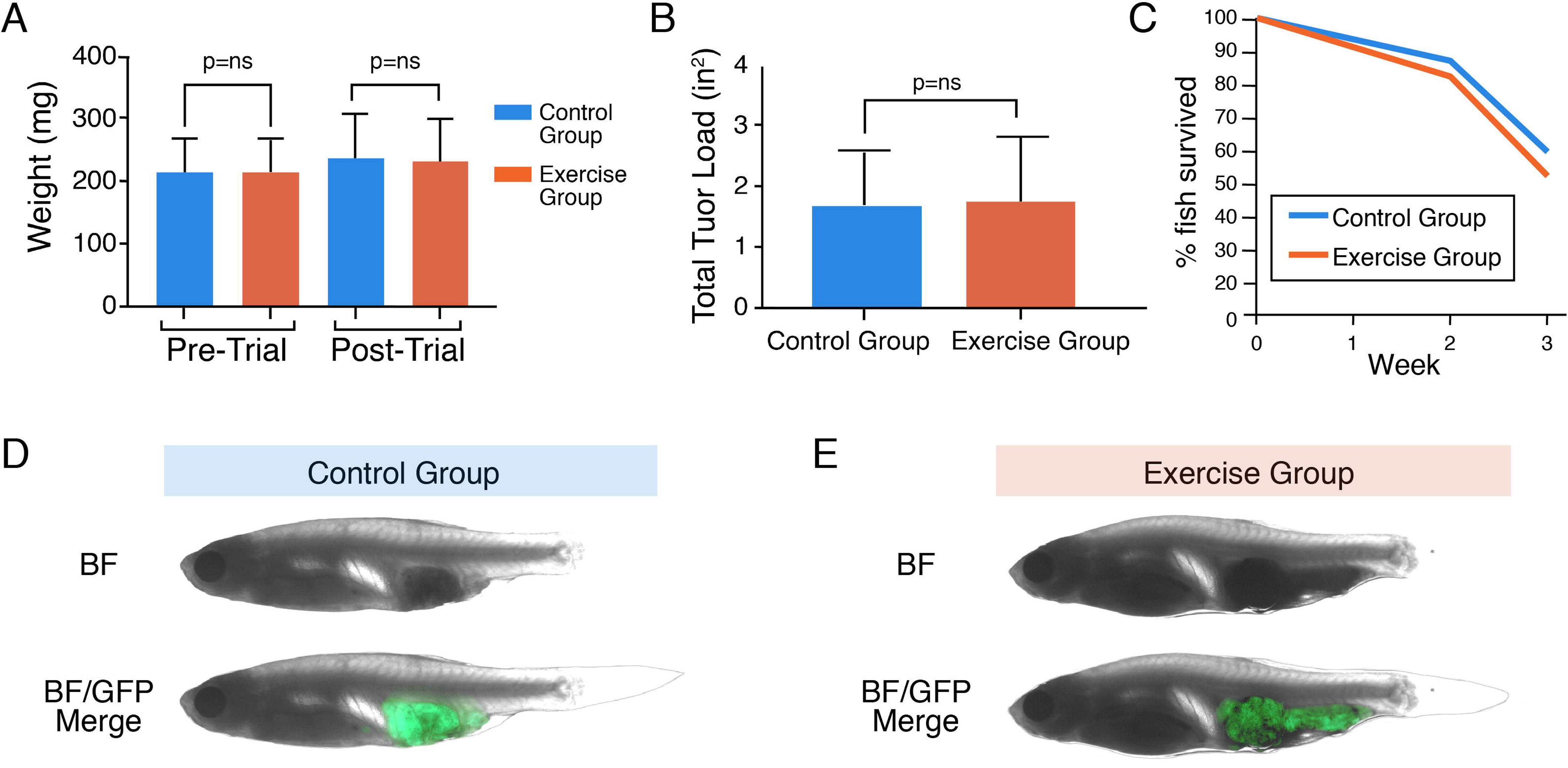
Adult exercise trial in melanoma. (A) Fish length over the period of the experiment showed no significant difference in control vs. exercise group. (B) Overall melanoma tumor size was assessed by measuring GFP+ area of the ZMEL1-GFP cells, after correcting for background autofluorescence. There were no significant differences between the control and exercise groups in this assay. (p=NS, Mann Whitney test). (C) Overall survival was similar between control and exercised fish. (D) Representative fish are shown at week 1 and 2 of the trial.

## Discussion

In this proof of principle study, we have established the zebrafish as a model to investigate exercise regulation of cancer. Although numerous studies have utilized the zebrafish as a model for cancer biology, and several as a model in exercise physiology (Massé et al., 2013; Wang et al., 2011; Conradsen and McGuigan, 2015; Gilbert et al., 2014), to our knowledge none have addressed whether these could be combined to specifically study the exercise - tumorigenesis link. Our data demonstrates that in at least one of our assays, melanoma larval transplantation, endurance exercise significantly inhibits melanoma growth *in vivo.*

The majority of studies addressing the mechanisms of exercise in cancer have focused on rodent models (Ashcraft et al., 2016). These are powerful for addressing fundamental issues of mammalian biology, although rodents are difficult to use for performing “unbiased” screens to identify unexpected pathways whereby exercise may regulate cancer. Such unbiased screening approaches require very large numbers of animals, a limiting factor in most rodent studies. In contrast, the zebrafish offers a major advantage in this regard since hundreds to thousands of animals can be studied simultaneously, providing appropriate statistical power to detect small differences.

It is increasingly appreciated that many cancer phenotypes are dependent upon interactions of tumor cells with their surrounding microenvironment (Quail and Joyce, 2013). Less well appreciated is how tumors cells interact with their “macroenvironment”, or the parts of the body very far away from the tumor itself. These interactions are especially important in exercise physiology, where virtually all organ systems can be affected, and lead to profound humoral and hormonal changes (Koelwyn et al., 2017). Factors such as insulin and IGF, which are strongly affected by exercise and impact liver and muscle, exert equally powerful effects on cancer cells (Capoluongo, 2011). Because these tumor-environment interactions are dynamic in time and space, dissecting these complex interactions in vivo is perhaps where the zebrafish will be best utilized. We and others have shown that the fish can be used to genetically or chemically interrogate both tumor cell as well as microenvironmental factors (Fior et al., 2017; Roh-Johnson et al., 2017). For example, we recently demonstrated using CRISPRs that knockout of microenvironmental endothelins strongly abrogates melanoma growth and metastasis (Kim et al., 2017). We envision using this CRISPR approach on a much larger scale, which will offer us the opportunity to functionally probe how tumor cells interact with their micro/macroenvironment during endurance exercise.

Another potential manner in which the zebrafish can be used to probe exercise oncology is through small molecule screens (Tamplin et al., 2012; North et al., 2007; Kaufman et al., 2009). In these screens, thousands of zebrafish embryos/larvae are exposed to a wide range of small molecules and a relevant phenotype is read out. We and others have used these approaches to identify small molecule modulators of neural crest development (White et al., 2011), BMP signaling (Yu et al., 2008), and factors affecting anxiety (Lundegaard et al., 2015). This technology was one of our primary reasons for testing the melanoma/exercise interaction using the ZEST system, since it opens up the possibility of finding small molecules that modulate how cancer cells respond to exercise. Such molecules may have direct therapeutic implications.

One unexpected finding from our study was that exercise only affected tumor growth in the larval, but not adult, assay. There are several major differences between these models. First, the larval fish are much younger and still rapidly developing. Although this larval model has been used for many years as a readout of cancer growth, it is possible that the younger fish are especially sensitive to the effects of exercise due to a competition for resources between the developing larvae versus the cancer cells. Second, the adult animals are irradiated before tumor implantation in order to suppress the immune system to prevent MHC mismatch between the ZMEL1-GFP cells and the recipient casper fish (which come from different strains). This immunosuppression is not necessary in the larval animal since they have innate NK cell immunity but not adaptive T-cell immunity at this age. In line with this, mouse studies have shown that exercise has a protective effect after implantation of B16 melanoma cells or use of transgenic Tg(Grm1)EPv melanoma models, an effect that is partly mediated by NK cell mobilization (Pedersen et al., 2016). One area for future exploration is to apply our exercise methods to transgenic zebrafish models (as opposed to transplantation models) since they will have a fully intact repertoire of both innate and adaptive immune cells, and then selectively ablate subpopulations of immune cells (i.e. NK cells).

While the zebrafish has clear advantages in terms of genetics and imaging, we recognize that there are important differences between exercise in fish versus mammals. One key difference is temperature - fish grow at 28C whereas mammals grow at 37C. Related to this, because fish are poikilothermic and do not regulate their temperature by sweating, they are subject to much greater swings in temperature than mammals that internally regulate this. Another key difference is that the fish are not subject to gravitational effects on skeletal muscle in the same manner as mammals, nor do they have a bony skeleton which is weight bearing. This will limit the amount of mechanical loading that their muscles can experience during prolonged exercise. Finally, and not least important, far less is known about the interplay between the various cell types affected in exercise. For example, the relationship between adipocytes, skeletal muscle and the immune system are largely unknown in this system. Given the increasing recognition that the immune system plays an intimate role in the response to exercise (Pedersen et al., 2016), this will be an area that requires further development. Finally, one question our study will help to address is to what extent exercise affects specific cancer types. There are now a multitude of cancer models in the zebrafish, including melanoma, pancreatic tumors, hematopoietic tumors, sarcoma and others. When used with our exercise system, each of these cancer models can now be efficiently studied to dissect the complex manner in which exercise affects each individual tumor type.

## Materials and Methods

### Cell Culture

The ZMEL1 cell line is derived from a primary zebrafish melanoma tumor overexpressing eGFP in a BRAF^V600E^ and p53^−/-^ background. The ZMEL1 cells were cultured in DMEM with 10% FBS, 1% Glutamax, and 1% penicillin-streptomycin and kept in an incubator at 28°C. In preparation for transplantation, ZMEL1 were trypsinized, spun down, and re-suspended in 90% 1xPBS and 10% sterile water.

### Fish Husbandry

All zebrafish were housed in a temperature (28.5°C) and light-controlled (14h on, 10h off) room. Fish were housed at a density of 5-10 fish per liter, and fed 3 times per day using brine shrimp and pelleted zebrafish food. All anesthesia was done using Tricaine (Western Chemical Incorporated) with a stock of 4g/L (protected for light) and diluted until the fish was immobilized. All procedures adhered to IACUC protocol #12-05-008 through Memorial Sloan Kettering Cancer Center.

### Adult Transplant

Recipient adult fish between the age of 2 months post fertilization and 12 months post fertilization were irradiated with 20 Gy of total gamma irradiation over the course of two days (10Gy/day). Irradiation of the adult recipient is necessary due to a MHC mismatch between the ZMEL1 cell line and the casper recipient. Irradiated fish were housed in their tanks for 2 days, during which feeding was held. Fish were transplanted on the third day post irradiation. ZMEL1 cells at about 90% confluency were trypsinized, spun down, and re-suspended at a density of about 167,000 cells/microliter of 1xPBS. Prior to transplant, fish were anesthetized with Tricaine and then placed on a damp towel. Approximately 500,000 ZMEL1 cells suspended in 3 microliters of 1xPBS were transplanted subcutaneously into the ventral flank of each irradiated adult fish using a Hamilton syringe (Hamilton 701N syringe, 10 microliters, 26s ga). The tumors engrafted and grew over a period of 14 days after which they were imaged.

### Larval Transplant

ZMEL1 cells at about 90% confluency were trypsinized, spun down, and re-suspended in 90% 1xPBS and 10% distilled water at a density of 25 cells/microliter. Larval zebrafish at around 48 hrs. post fertilization were anesthetized with Tricaine and transplanted with around 25 ZMEL1 cells suspended in 1 microliter using a borosilicate glass capillary needle attached to a microinjection apparatus. Cells were transplanted directly into the yolk sac of the larval embryo. The embryo recipients at 48 hours post fertilization do not require any preconditioning radiation since they do not yet have a mature adaptive immune system.

### Adult Swim Exercise

Adult zebrafish were exercised using the Loligo swim tunnel respirometer 5L (120V/60Hz). Flow was measured using a handheld digital flow meter (Loligo Systems). Flow velocity readings were taken at 6 locations within the swim tunnel and averaged to calculate the dose of exercise. Before each exercise session, water from the tank was replaced with fresh system water. The exercise fish group was placed in the flow chamber with no flow and acclimated for 5 minutes. The velocity of flow was increased steadily over a period of 5 minutes until the final velocity was reached. Adult fish were exercised at a constant speed of 25 cm/s for 3 hrs./day, 5 days/week. The control group remained in the flow chamber for 3 hrs/day and 5 days/week with no water flow.

### Embryo Swim Exercise

Embryonic zebrafish were placed into a custom built zebrafish embryo swim tunnel (ZEST) made from acrylic round tube (7/8” OD, 5/8” ID, McMaster-Carr). Each end of the tube was covered with moisture-resistant polyester mesh (.0148” opening, McMaster-Carr). One end of the tube was made removable to allow for the addition and removal of embryos from the swim tunnel (Figure 1). The ZEST was placed into the swim chamber of the Loligo swim tunnel respirometer 5L (120V/60Hz). Flow was measured using a handheld digital flow meter (Loligo Systems). Flow velocity readings were taken at 6 locations within the swim tunnel and averaged to calculate the dose of exercise. Before each exercise session, water from the tank was replaced with fresh system water. The exercise group was placed in the flow chamber with no flow and acclimated for 5 minutes. Embryonic zebrafish were transferred into the ZEST, which was placed into the swim chamber of the 5L swim tunnel respirometer and allowed to acclimate for 5 minutes. The speed of the flow was increased steadily over a period of 5 minutes until a final velocity of 8 cm/s was reached. The embryos in the exercise group were maintained at a speed of 8cm/s for 6 hrs./day, 5 days/week. The control group was transferred into the ZEST which was placed into the swim chamber of the 5L swim tunnel respirometer where they sat for 6 hrs./day, 5 days/week with no water flow.

### Imaging and Image Processing

All fish were anesthetized with Tricaine and placed onto an agar coated petri dish. The fish were imaged from above using a Zeiss Axio Zoom V16 Fluorescence Stereo Zoom Microscope with a 0.6x or 1.6X lens. Each fish was successively imaged using brightfield, GFP and Rhodamine filter sets on both sides. The exposure times for each group were determined at day 1 and kept fixed throughout the entire experiment. Raw image files (CZI) for each fish were then exported using Zen into high resolution TIFFs which could then be used for downstream image analysis in Image J.

### Downstream image analysis in Image J

High resolution TIFFs were imported into Image J. For analysis of the adult fish, only the GFP channel was imported. Total GFP was quantified for each image. For analysis of the embryonic zebrafish, the GFP channel and Rhodamine channel were both imported. The Rhodamine channel was used as a background subtraction to remove autofluorescence. After the background subtraction, total GFP was quantified.

## Acknowledgments

The authors would like to thank Wenjing Wu for assistance with illustrations.

R.M.W. is supported by the NIH Director’s New Innovator Award (DP2CA186572), Mentored Clinical Scientist Research Career Development Award (K08AR055368), the Melanoma Research Alliance, The Pershing Square Sohn Foundation, The Alan and Sandra Gerry Metastasis Research Initiative at the Memorial Sloan Kettering Cancer Center, The Harry J. Lloyd Foundation and Consano. L.W.J. is supported by research grants from the National Cancer Institute (5R01CA179992), AKTIV Against Cancer and the Memorial Sloan Kettering Cancer Center Support Grant/Core Grant (P30CA008748). N.R.C. was supported by a Medical Scientist Training Program grant from the National Institute of General Medical Sciences of the National Institutes of Health under award number T32GM007739 to the Weill Cornell/Rockefeller/Sloan-Kettering Tri-Institutional MD-PhD Program

## Author Contributions

R.M.W. and L.J. conceived the study. A.Y. performed all zebrafish experiments. N.C. assisted with image analysis.

